# ETV4 is necessary for estrogen signaling and growth in endometrial cancer cells

**DOI:** 10.1101/617142

**Authors:** Adriana C. Rodriguez, Jeffery M. Vahrenkamp, Kristofer C. Berrett, Kathleen A. Clark, Katrin P. Guillen, Sandra D. Scherer, Chieh-Hsiang Yang, Bryan E. Welm, Margit M. Janát-Amsbury, Barbara J. Graves, Jason Gertz

## Abstract

Estrogen signaling through estrogen receptor alpha (ER) plays a major role in endometrial cancer risk and progression; however, the molecular mechanisms underlying ER’s regulatory role in endometrial cancer are poorly understood. In breast cancer cells, ER genomic binding is enabled by FOXA1 and GATA3, but the transcription factors that control ER genomic binding in endometrial cancer cells remain unknown. We previously identified ETV4 as a candidate factor controlling ER genomic binding in endometrial cancer cells and here we explore the functional importance of ETV4. Homozygous deletion of *ETV4*, using CRISPR/Cas9, led to greatly reduced ER binding at the majority of loci normally bound by ER. Consistent with the dramatic loss of ER binding, the gene expression response to estradiol was dampened for most genes. ETV4 contributes to estrogen signaling in two distinct ways; ETV4 loss impacts chromatin accessibility at some ER bound loci and impairs ER nuclear translocation. The diminished estrogen signaling upon *ETV4* deletion led to decreased growth, particularly in 3D culture where hollow organoids were formed and *in vivo* in the context of estrogen dependent growth. Our results show that ETV4 plays an important role in estrogen signaling in endometrial cancer cells.

## INTRODUCTION

Endometrial cancer is the fourth most common cancer among women and the most common gynecological cancer, leading to over 10,000 deaths in the United States each year[1]. Type 1 endometrial cancer makes up ~85% of cases, and the vast majority of these cases express estrogen receptor ɑ (ER), a steroid hormone receptor[2]. Risk factors such as unopposed estrogen therapy, tamoxifen treatment for breast cancer, lifetime exposure to estrogens as captured by the number of fertility cycles, and obesity, which leads to peripheral estrogen production, all indicate a central role for estrogen signaling in endometrial cancer development (reviewed in [3]). Endometrial cancer mutations in the ligand binding domain of ER[4, 5], which likely confer constitutive activity, implicate ER as the main mediator of estrogen signaling in endometrial tumors. Endometrial cancer is most commonly treated by hysterectomy; however, progestins, synthetic progesterone, serve as an effective alternative treatment[6]. Progestins bind to and activate progesterone receptor, which antagonizes the mitogenic effects of ER[7]. These risk factors and treatment strategies for endometrial cancer point to ER as a key oncogene in the disease.

While ER plays an important role in endometrial cancer, little is understood about ER’s molecular actions in endometrial cancer cells. Many of the molecular details concerning ER have come from extensive work in breast cancer[8–15], where ER has a critical oncogenic role. These findings include regulatory co-factors that ER utilizes to regulate gene expression[16–18]. In addition, FOXA1 and GATA3 have been shown to be necessary pioneer factors that maintain accessible chromatin for ER in breast cancer cells[19–21]. ER binds to the genome in a cell type-specific manner with disparate binding between breast and endometrial cancer cells[14, 22]; therefore, different transcription factors are likely to control ER binding in endometrial cancer cells. In concordance with this idea, *FOXA1* and *GATA3* exhibit low expression in endometrial tumors[4]. As ER has a different set of genomic binding sites and FOXA1 only overlaps with ~8% of ER’s binding sites in endometrial cancer cells and tumors, compared to 57% in breast cancer cells[14, 22], it is likely that different transcription factors are responsible for controlling ER’s genomic interactions in endometrial cancer cells.

In a previous study, we identified ETV4, an ETS factor, as playing a potential role in endometrial cancer-specific ER binding[14]. *De novo* motif discovery of ER binding sites specific to endometrial cancer cells identified the ETS factor motif. Of the ETS factors, ETV4 is specifically expressed in the endometrial cancer cell line, Ishikawa, as opposed to the ER-positive breast cancer cell line, T-47D. ChIP-seq in Ishikawa cells revealed that ETV4 overlaps with 45% of ER bound sites[14]; however, the functional importance of this overlap has not been tested. The potential role of ETV4 in controlling ER binding in endometrial cancer is an intriguing possibility, as other members of the ETS family have been shown to have pioneering capabilities for steroid hormone receptors. In prostate cancer, overexpression of ETV1, an ETS factor closely related to ETV4, has been shown to reprogram androgen receptor genome binding and associated transcription regulation, leading to the formation of prostatic intraepithelial neoplasia[23], and contributing to more aggressive tumors[24]. Similar effects have been shown for ERG, another ETS factor commonly overexpressed in prostate cancer[25]. The mechanisms underlying these effects on androgen receptor are still unclear; however, a physical interaction has been observed between ETV1 and androgen receptor[23]. Though ETS factors in prostate cancer are being widely studied, the functional role of ETV4 in ER’s genomic binding in endometrial cancer has not been evaluated.

In this study, we use CRISPR/Cas9 to knockout *ETV4* in Ishikawa cells in order to determine the importance of ETV4 in controlling ER genomic binding. Despite having no effect on ER protein levels, *ETV4* knockout led to greatly reduced ER binding, with the majority of sites exhibiting a significant reduction in ChIP-seq signal. Consistent with the reduction in ER binding, the transcriptional response to 17β-estradiol (E2) was diminished due to *ETV4* knockout. Chromatin accessibility was also reduced, but only at a subset of sites suggesting a minimal pioneering role for ETV4 in endometrial cancer cells. Unexpectedly, we also discovered an increase in cytoplasmic ER, suggesting that ETV4 contributes to ER activation. Genetic deletion of *ETV4* caused phenotypic changes, including a reduction in growth that is intensified in 3D culture. Overall, we find that *ETV4* is necessary for genomic binding of ER, gene regulation by E2, and growth, in endometrial cancer cells.

## RESULTS

### ETS motif is found at endometrial cancer ER binding sites and *ETV4* is overexpressed in endometrial tumors

Previous studies have shown that ER binds to different genomic loci in breast and endometrial cancer cell lines[14] and tumors[22]. To gain a more comprehensive view of ER binding across breast and endometrial tumors, we performed ChIP-seq to map ER genomic binding in 5 endometrial cancer xenografts (see Table S1 for clinical information) and combined the data with published ER ChIP-seq datasets[10, 22], representing 9 breast tumors and 10 endometrial tumors in total. Unsupervised clustering distinctly segregated the samples by cancer type (Fig. 1A), with the Ishikawa endometrial cancer cell line grouping closely to endometrial cancer patient samples. Previous *de novo* motif discovery at Ishikawa specific ER binding sites found enrichment of the ETS motif[14]. Analysis of ER binding sites identified in tumors revealed a similar enrichment of an ETS motif (Fig. 1B), with a 2.9-fold enrichment compared to shuffled sequence controls. Droog et al.[22] identified the FOXA binding motif at ER bound loci in endometrial tumors. We found that the ETS motif was more enriched than the FOXA motif (1.6-fold enrichment over shuffled sequences) at ER bound sites in endometrial tumors (p-value < 0.0001, t-test). Additionally, the enrichment for the FOXA motif was significantly higher in breast tumor ER binding sites (p-value = 0.0094, t-test, Fig. 1C), indicating that forkhead factors (e.g. FOXA1) may play a bigger role in breast cancer ER genomic binding site selection while ETS factors may play a bigger role in endometrial cancer ER binding site selection.

**Figure 1.**
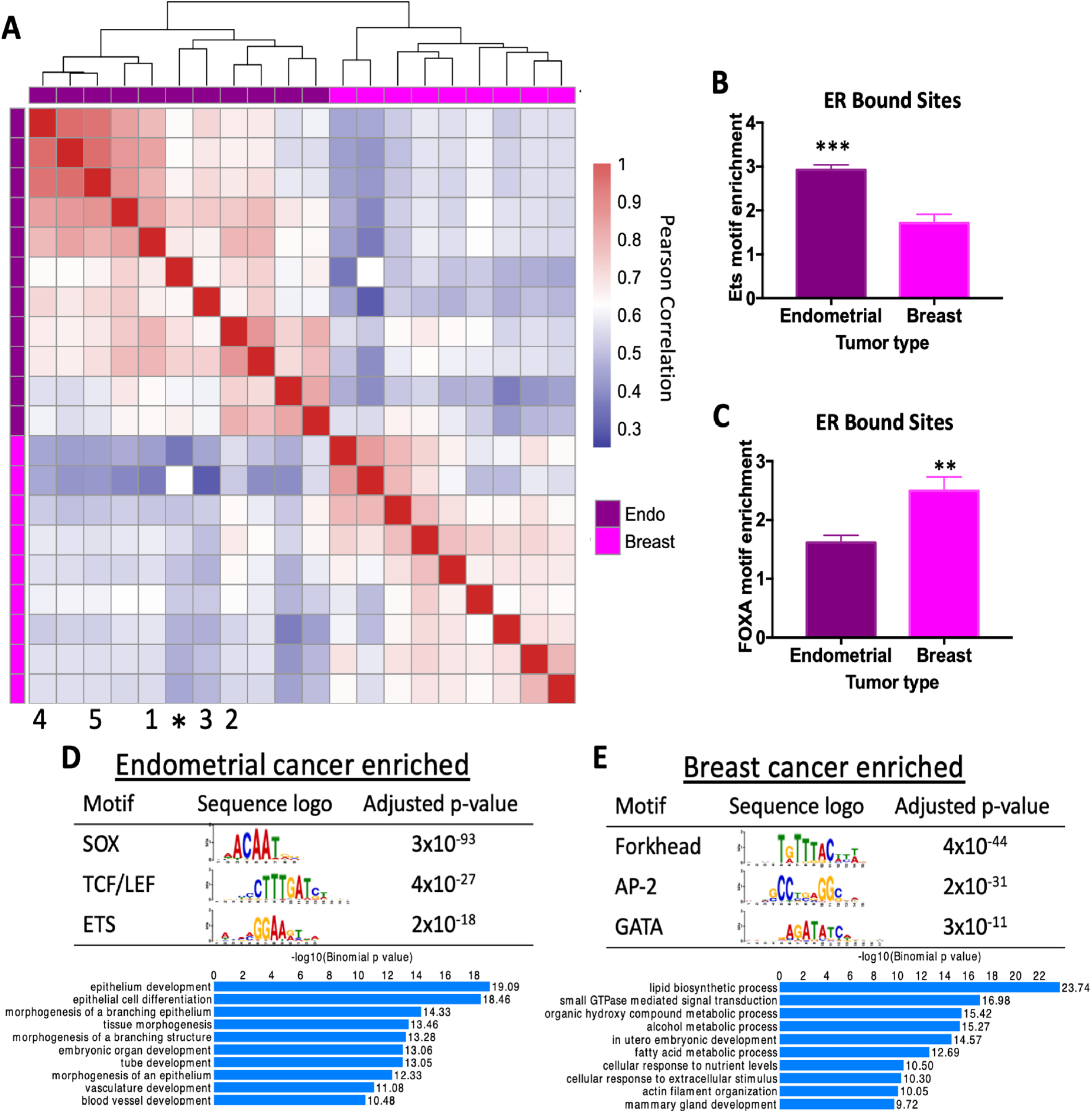
ER binding and associated transcription factor motifs are cancer type-specific. A) Heatmap comparing correlation in ER genome binding in endometrial cancer and breast cancer patient samples shows that ER binding patterns are specific to cancer type. * represents Ishikawa endometrial cancer cell line. Numbers indicate PDX model (1 = EC-PDX-004, 2 = EC-PDX-005, 3 = EC-PDX-018, 4 = EC-PDX-021, and 5 = EC-PDX-026). Sequence analysis of ER ChIP-seq data shows ETS motif (p-value=0.0003, unpaired t-test) (B) and forkhead factor motif (p-value= 0.0094, unpaired t-test) (C) enrichment in ER bound sites in endometrial tumors and breast tumors. The top motifs from differential motif analysis and the top GO biological processes for nearby genes are shown for endometrial cancer-specific ER bound sites (D) and breast cancer-specific ER bound sites (E).

To complement this directed motif analysis, we searched for motifs enriched in breast-specific ER bound sites and endometrial cancer-specific ER bound sites, defined as being present in at least 30% of one cancer type’s samples and none of the other cancer type’s samples. Using analysis of motif enrichment from the MEME suite[26], we found a forkhead motif, AP-2 motif, and GATA motif, enriched in breast cancer-specific ER sites (Fig. 1E). The importance of FOXA1 and GATA3 in ER genomic binding in breast cancer cells[19, 27] corroborates this finding. In endometrial cancer-specific sites, we found the ETS motif as expected in addition to SOX motifs and TCF/LEF motifs (Fig. 1D), indicating that several factors may be involved in ER genomic binding that are endometrial cancer-specific. Endometrial cancer-specific and breast cancer-specific ER bound sites were also found nearby different sets of genes (Fig. 1D,E) based on GREAT analysis [28]. Genes near the endometrial cancer-specific ER bound sites were enriched in epithelial development and branching morphogenesis, while genes nearby breast cancer-specific ER bound sites were enriched in lipid biosynthesis and regulation of small GTPases.

In analyzing the expression levels of ETS family members, we previously identified *ETV4* as having high expression in Ishikawa cells compared to T-47D, a breast cancer cell line. We also found that ETV4 bound to ~45% of Ishikawa-specific ER binding sites, indicating that ETV4 may be contributing to endometrial cancer-specific ER binding[14]. Using TCGA data, we found that while *ETV4* levels are low in both breast cancer and normal breast tissue, there is an average 3.3-fold increase in *ETV4* expression in endometrial cancer samples compared to normal endometrial tissue (p-value = 1×10^−21^ [29], Fig. S1A)[30]. Focusing specifically on endometrioid histology endometrial tumors, we found that ETV4 and ER are co-expressed at high levels in the majority of tumors (Fig. S1B). Analysis of the endometrial cancer PDX models revealed high ETV4 protein expression and nuclear staining in three of the five models (Fig. S1C). Based on these observations, we hypothesize that ETV4 is important in ER genomic binding in endometrial cancer cells and we therefore sought to functionally dissect ETV4’s role in estrogen signaling in endometrial cancer.

### *ETV4* loss substantially reduces ER genomic binding

To assess if ETV4 is involved in regulating ER genomic binding, we knocked out *ETV4* in Ishikawa cells using CRISPR/Cas9 with homology directed repair plasmids (see Methods). We isolated individual clones and validated two clones by ETV4 Western blot and qPCR (Fig. 2A, Fig. S2). The expression of ER was unaffected by *ETV4* knockout (KO) (Fig. 2B). We mapped ER binding using ChIP-seq, after a 1-hour induction with 10nM E2, and identified 6,263 binding sites in wildtype Ishikawa cells. ER genomic binding was largely reduced in both *ETV4* KO clones, with only 1,957 sites (69% reduction) detected in one clone and 1,256 (80% reduction) detected in the other (Fig. 2C). The reduction in ER binding was confirmed with a second biological replicate and 69% of ER bound sites exhibited at least a 50% reduction in signal intensity on average (Fig. 2D, example loci are shown in Fig. 2E). Since these ChIP-seq experiments were performed using an antibody that is no longer commercially available, we repeated the experiment using a monoclonal ER antibody and found a similar reduction. While many more peaks were called with the monoclonal antibody, a 42% reduction in ER binding was observed (139,036 wildtype peaks vs. 81,232 ETV4 knockout peaks). To determine if the reduction in ER genomic binding is due to long-term loss of ETV4 or more immediate effects, we performed siRNA knockdown of ETV4 in wildtype Ishikawa cells. We found that protein levels of ETV4 were stable 72 hours post-transfection but lost 96 hours post-transfection (Fig. S3A,B). ER genomic binding was relatively unchanged 72 hours post-transfection (16,034 vs 16,545 ER bound sites in non-targeting controls and ETV4 siRNA). At 96 hours post-transfection, ER binding was reduced from 18,266 to 5,565 (70% reduction) with ETV4 siRNA (Fig. S3C). These findings indicate that ETV4 loss has immediate effects on ER genomic binding. The overall large reduction in ER genomic binding with *ETV4* loss is unexpected given that ETV4 bound sites overlap with only 45% of Ishikawa-specific ER bound sites; however, a similar pattern was observed with FOXA1 knockdown in MCF-7 cells, a breast cancer cell line[19].

**Figure 2.**
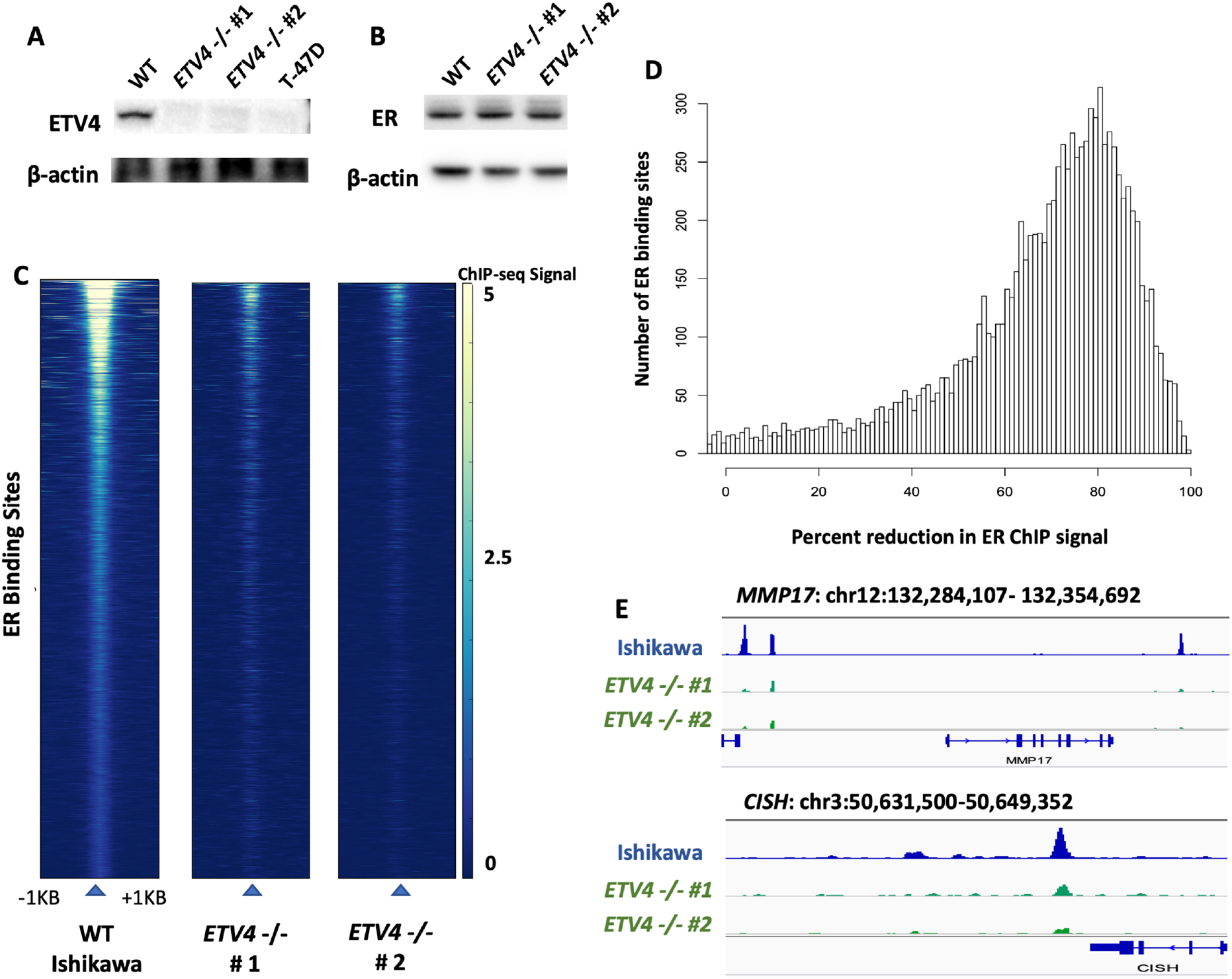
Genetic deletion of *ETV4* leads to a reduction of ER genomic binding. Western blots show protein expression of ETV4 (A) and ER (B) in wildtype and ETV4 knockout Ishikawa cells. Absence of ETV4 protein in two knockout clones and T-47D (negative control) is indicated by Western blot. C) Heatmaps of ER ChIP-seq signal show greatly reduced ER binding in both of the knockout clones. D) Histogram displays the loss of ER ChIP-seq signal in ETV4 knockout clones, data represents two clones and two biological replicates per clone. E) Examples of decreased ER ChIP-seq signal at *MMP17* and *CISH*, two known estrogen induced genes in Ishikawa cells[31], are shown.

To validate that the reduction in ER genomic binding is due to the loss of *ETV4*, we rescued expression of *ETV4* in KO clones, and overexpressed *ETV4* in wildtype cells for comparison using the same expression construct (Fig. S4). ChIP-seq of the rescue lines shows a restoration of ER binding, with a 35% loss of ER binding compared to wildtype, instead of the 69% loss observed in *ETV4* KO lines (Fig. S4). We also observed a greater than two-fold increase in ER genomic binding between wildtype cells and wildtype cells with *ETV4* overexpression while maintaining a high level of overlap (Fig. S4). The results support the idea that levels of ETV4 are correlated with the amount of ER genomic binding.

One possible explanation for the sites that remain bound after *ETV4* KO, is that these sites represent direct ER binding sites with full palindromic estrogen response elements (EREs). Of the sites that are still bound by ER in the ETV4 KO clones, 23% harbored full ERE matches. This is significantly higher than sites that were lost upon ETV4 KO (14%, p-value < 2.2×10^−16^, Fisher’s exact test). *De novo* motif discovery in the sites lost upon knockout show enrichment of the ETS motif, as expected. In agreement with this observation, loci with over 50% loss in ER ChIP-seq signal upon ETV4 loss were 2.4-fold more likely to overlap with ETV4 than loci that did not exhibit 50% loss in signal (p-value < 2.2×10^−16^, Fisher’s Exact Test). These results suggest that ETV4 is necessary for a large portion of ER genomic binding in endometrial cancer cells; however, sites with full EREs and no ETV4 binding are less impacted.

### ETV4 loss diminishes the transcriptional response to estradiol

In endometrial cancer cells, ER bound by estrogens leads to an induction of transcription at hundreds of genes. To test whether the loss of *ETV4*, and the subsequent loss of ER binding, leads to a reduction in the transcriptional response to estrogens, we performed RNA-seq on wildtype and ETV4 KO lines grown in hormone depleted media and treated for 8 hours with vehicle (DMSO) or E2. The most estrogen responsive genes were dramatically affected by *ETV4* KO, with 9 of the top 15 E2-induced genes showing a significantly reduced E2 response in *ETV4* KO cells (Fig. 3A). The estrogen response was also diminished genome-wide. Fig. 3B shows a comparison of the E2 response between wildtype and *ETV4* KO cells, where many estrogen induced genes exhibit a reduced (off diagonal) response in the *ETV4* KO cells. The reduced E2 response was observed for both up- and down-regulated genes indicating that ETV4 loss does not only impact gene activation through ER. As a control, the two KO clones show no distinct trend towards either axis, with the majority of genes having similar estrogen responses in both lines (Fig. 3C). Of the 3,980 genes that gain expression upon estrogen induction in wildtype Ishikawa cells, 3,412 (85%) are more highly induced in wildtype cells than knockouts, while only 568 (15%) are more highly induced in *ETV4* KO cells (p-value < 2.2×10^−16^ Wilcoxon test comparing fold changes)(Fig. S5A). To examine the relationship between ETV4 genomic binding and genes that lose or maintain estrogen inductions, we calculated the distance between each gene’s transcription start site and the nearest ETV4 bound site in wildtype Ishikawa cells. Genes that lose estrogen induction are significantly closer to ETV4 bound sites than genes that maintain an estrogen induction (p-value = 0.00022, Wilcoxon test, Fig. S5B). Overall, these results suggest that loss of *ETV4* largely reduces the transcriptional response to estrogens in endometrial cancer cells.

**Figure 3.**
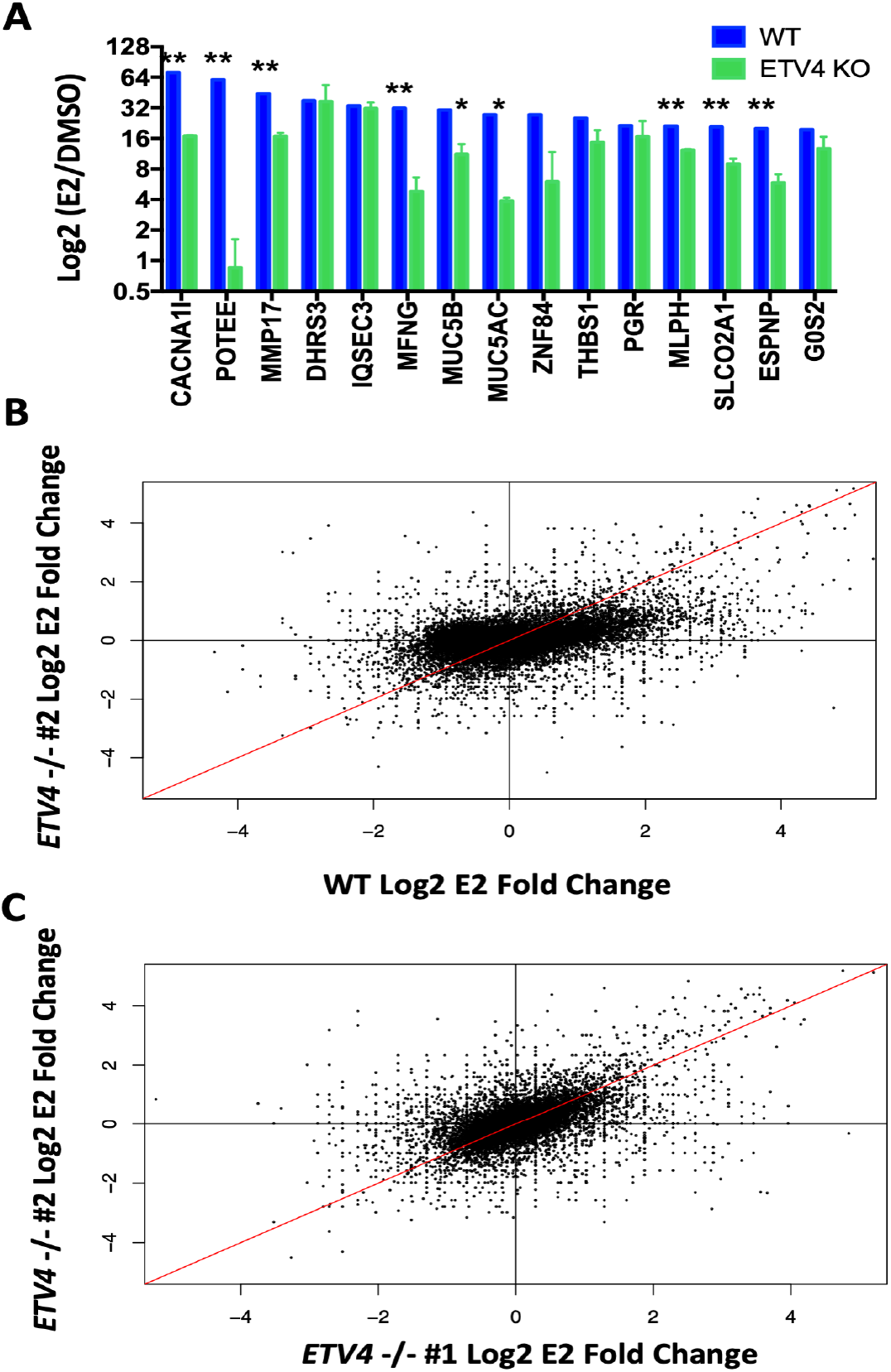
ETV4 is necessary for estrogen regulated gene expression. A) The top estrogen-induced genes in wildtype Ishikawa cells (blue) have significantly reduced (**p-value > 0.01; *p-value < 0.05) estrogen induction in *ETV4* KO clones as tested by RNA-seq. B) Comparison of fold changes in estrogen induced expression of all genes shows reduced estrogen response globally in knockout (y-axis) compared to wildtype (x-axis). C) Comparison of both knockout clones shows they share similar estrogen response patterns.

### Chromatin accessibility is impacted by ETV4 loss at some ER binding sites

One potential explanation for how ETV4 is contributing to ER genomic binding in endometrial cancer cells is that ETV4 acts as a pioneer factor and maintains accessible chromatin at ER binding sites. In order to determine how ETV4 impacts chromatin accessibility, we performed Assay for Transposase Accessible Chromatin followed by sequencing (ATAC-seq)[32] in wildtype and *ETV4* KO cells. We did not observe large genome-wide changes in chromatin accessibility due to *ETV4* loss (Fig. S6); however, some of the loci bound by both ETV4 and ER were impacted. We analyzed the 3,114 ATAC-seq sites that overlapped with ER and ETV4 binding and our analysis showed that 392 sites (13%) significantly lost chromatin accessibility in the knockout lines and 246 sites (8%) gained accessibility, both in an estrogen-independent manner (Fig. 4A). The observed changes in chromatin accessibility indicate that ETV4 is necessary to pioneer a small subset of sites, creating an accessible landscape for ER genomic binding; however, this effect does not explain the large-scale reduction in ER genomic binding with ETV4 loss.

**Figure 4.**
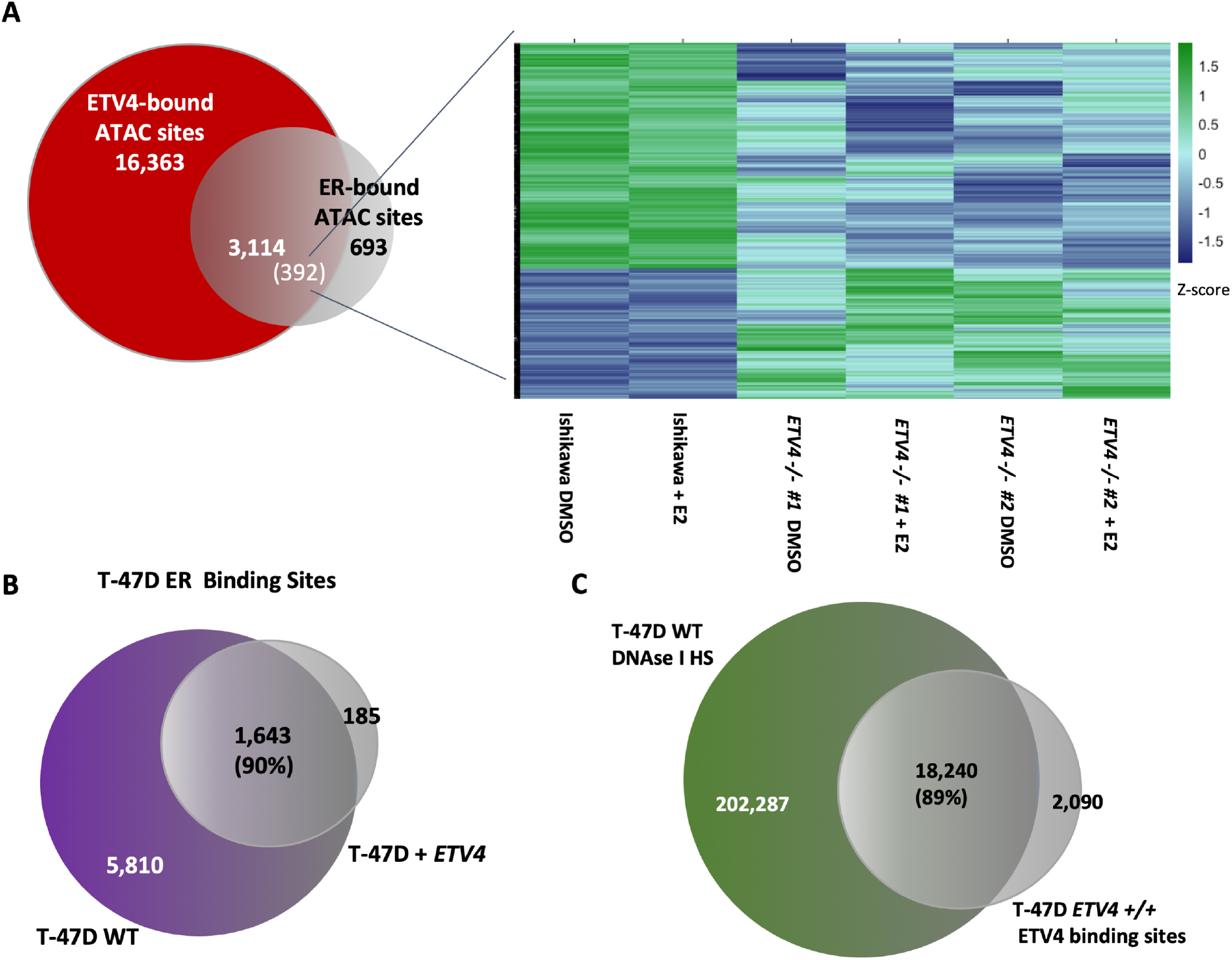
ETV4 only impacts chromatin accessibility at some ER bound sites and is not sufficient to reprogram ER binding in breast cancer cells. A) A subset of ATAC-seq sites bound by both ETV4 and ER in Ishikawa cells show significant changes in ATAC-seq signal. Heatmap shows ATAC-seq signal at significantly changing sites. B) The overlap in ER genomic binding between T-47D cells and T-47D cells with forced expression of *ETV4* is shown. C) Venn diagram shows the overlap between DNase I hypersensitive sites in wildtype T-47D cells and ChIP-seq of ETV4 (FLAG) in T-47D lines stably expressing *ETV4*, indicating that ectopically expressed ETV4 preferentially binds loci that are already accessible.

### *ETV4* is not sufficient to reprogram ER binding in a breast cancer cell line

In order to test the sufficiency of *ETV4* in controlling ER binding, we ectopically expressed FLAG-tagged *ETV4* in T-47D cells, an ER expressing breast cancer cell line (Fig. S7A). We first looked at ER genomic binding by ChIP-seq and compared the data with ER ChIP-seq in wildtype T-47D cells. ER binding site selection was not substantially changed by forced *ETV4* expression as 90% of ER binding sites overlapped between wildtype and ETV4 expressing T-47D cells (Fig. 4B, Fig. S7B). We next analyzed ETV4 binding in T-47D cells, using an antibody that recognizes FLAG, and found that ETV4 binding was substantially different in T-47D cells than Ishikawa cells with 54% overlap. Comparison of ETV4 genomic binding in T-47D with DNase I hypersensitivity sites in T-47D [14] revealed that 89% of the ETV4 binding sites are in regions of open chromatin (Fig. 4C). This data implies that ETV4 preferentially binds open chromatin instead of binding to a set of preferred sequences, and it is not sufficient to reprogram ER binding in another cell type. These findings together with the minimal changes in chromatin accessibility with *ETV4* loss are consistent with ETV4 not acting as a pioneer factor for ER.

### Loss of ETV4 leads to a change in ER localization

ETV4’s inability to dramatically impact chromatin accessibility implies it is affecting ER binding by another mechanism. ER is found both in the nucleus and cytoplasm in the absence of ligand and the distribution shifts towards the nucleus upon ligand binding [33, 34]. In order to test whether *ETV4* loss leads to decreased ER activation, we looked for a change of localization by isolating cytoplasmic and nuclear fractions from wildtype Ishikawa cells and the *ETV4* KO lines before analyzing ER levels (Fig. 5A). As expected, the ratio of nuclear to cytoplasmic ER increased upon a 1-hour E2 induction in wildtype Ishikawa cells from a relative ratio of 1 to 1.43 (p-value = 0.0389, unpaired t-test). The *ETV4* KO lines had a reduced ER nuclear to cytoplasmic relative ratio of only 0.434 (p-value = 0.0148, unpaired t-test, Fig. 5B) and the ratio remained low in the presence of E2, 0.359 (p-value= 0.0011, unpaired t-test, Fig. 5B). In the context of E2 treatment, we observed a 4-fold reduction in the ratio of nuclear to cytoplasmic ER due to ETV4 loss, which is similar to the 3.5-fold reduction observed in ER genomic binding. This significant reduction in ER’s nuclear to cytoplasmic ratio implies an ER shuttling defect associated with *ETV4* loss.

**Figure 5.**
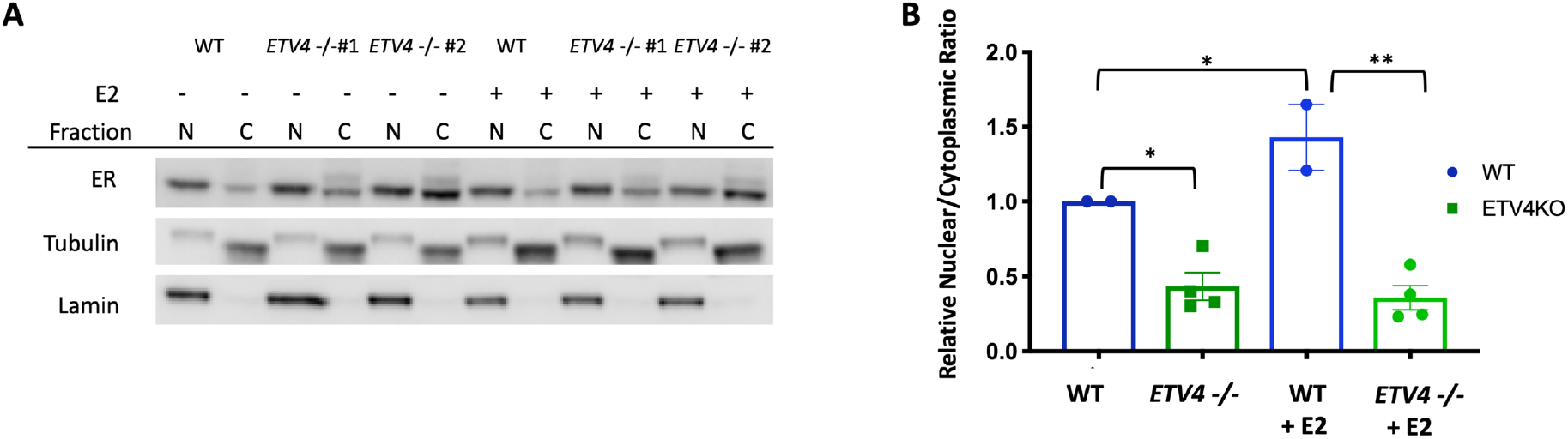
Cytoplasmic localization of ER is increased in the absence of *ETV4*. A) Western blot of ER levels in isolated nuclear (N) and cytoplasmic (C) fractions, after 1-hour treatments with vehicle (DMSO), or 10nM E2. Tubulin and lamin are shown as cytoplasmic and nuclear controls, respectively. B) Relative ratios of nuclear to cytoplasmic ER of duplicate experiments, normalized to the wildtype vehicle ratio, are shown. Error bars represent s.e.m. * = p-value < 0.05, ** indicates p-value < 0.005.

In order to determine whether this localization change is specific to ER or a general effect on steroid hormone receptors, the experiment was repeated for glucocorticoid receptor (GR). After a 1-hour induction with the GR ligand Dexamethasone (Dex), the ratio of nuclear GR to cytoplasmic GR increased 23-fold and in the *ETV4* KO lines a similar increase of 31-fold was observed (Fig. S8). The similar nuclear to cytoplasmic ratio of GR in the *ETV4* KO lines implies that the effect of *ETV4* loss on ER’s localization is specific to ER and not a general defect in steroid hormone receptor signaling.

### ETV4 knockout causes reduced growth in 3D culture

Since *ETV4* is required for the transcriptional response to estrogens, we hypothesized the *ETV4* KO would cause reduced growth in endometrial cancer cells that normally increase proliferation when exposed to E2. To explore this possibility, we performed growth assays both in 2D and 3D culture. Utilizing live cell imaging on the Incucyte ZOOM imaging system, we measured 2D culture growth rates. The cells were plated at low confluency in either full media (estrogenic), or hormone-depleted media and were imaged for 72 hours. We observed a 9% reduction in growth for *ETV4* KO cells in full serum media compared to wildtype cells (p-value = .007, unpaired t-test, Fig. S9). While in hormone-depleted media, all cells had similar growth rates. The reduced growth of *ETV4* KO cells in full serum media was similar to the growth rate of wildtype cells in hormone-depleted media (Fig. S9). This indicates that upon loss of *ETV4*, cell growth is reduced only in the presence of estrogens and other hormones.

We also analyzed growth in 3D culture. Cells were grown in matrigel in either full serum media or hormone-depleted media and ATP levels were measured as a proxy for growth. The results corroborated the findings from 2D culture, where *ETV4* KO significantly reduced growth only in full serum media (full media: p-value = 0.03, hormone-depleted media: p-value = 0.38, unpaired t-test, Fig. 6B). Image analysis of the *ETV4* KO organoids showed that many had a ring of live cells without any cells in the center (Fig. 6A). H&E staining revealed many hollow spheroids (Fig. 6C) and showed this phenotype to be present 2.3-fold more often in the knockout cells than in wildtype Ishikawa cells when grown in regular media (Fig. 6D). In hormone-depleted media, the phenotype is observed 1.7-fold more frequently in knockout cell spheroids than wildtype cell spheroids (Fig. 6E). The hollow spheroids do not appear to be caused by the death of internal cells, as very few cytox orange positive cells are observed (Fig. 6A). These observations indicate that in the absence of *ETV4*, cells are more likely to start forming structures with glandular morphology in 3D culture conditions that are reminiscent of the glandular structure found in the normal endometrium. In order to explore this further, we performed e-cadherin and pan-cytokeratin immunohistochemistry to look at markers of normal endometrium[35]. While e-cadherin levels remained negative in both the wildtype and the *ETV4* KO lines, pan-cytokeratin levels increased in the absence of *ETV4*, often lining the interior of the hollow organoids (Fig. S10). This pattern indicates a partial redifferentiation to normal endometrium with the loss of *ETV4*. Mucin staining using Periodic Acid Schiff (PAS) was negative, indicating any redifferentiation is more likely towards proliferative phase endometrium as opposed to secretory phase (Fig. S10). Overall, the loss of *ETV4* leads to a change in morphology that partially resembles a normal proliferative phase endometrium.

**Figure 6.**
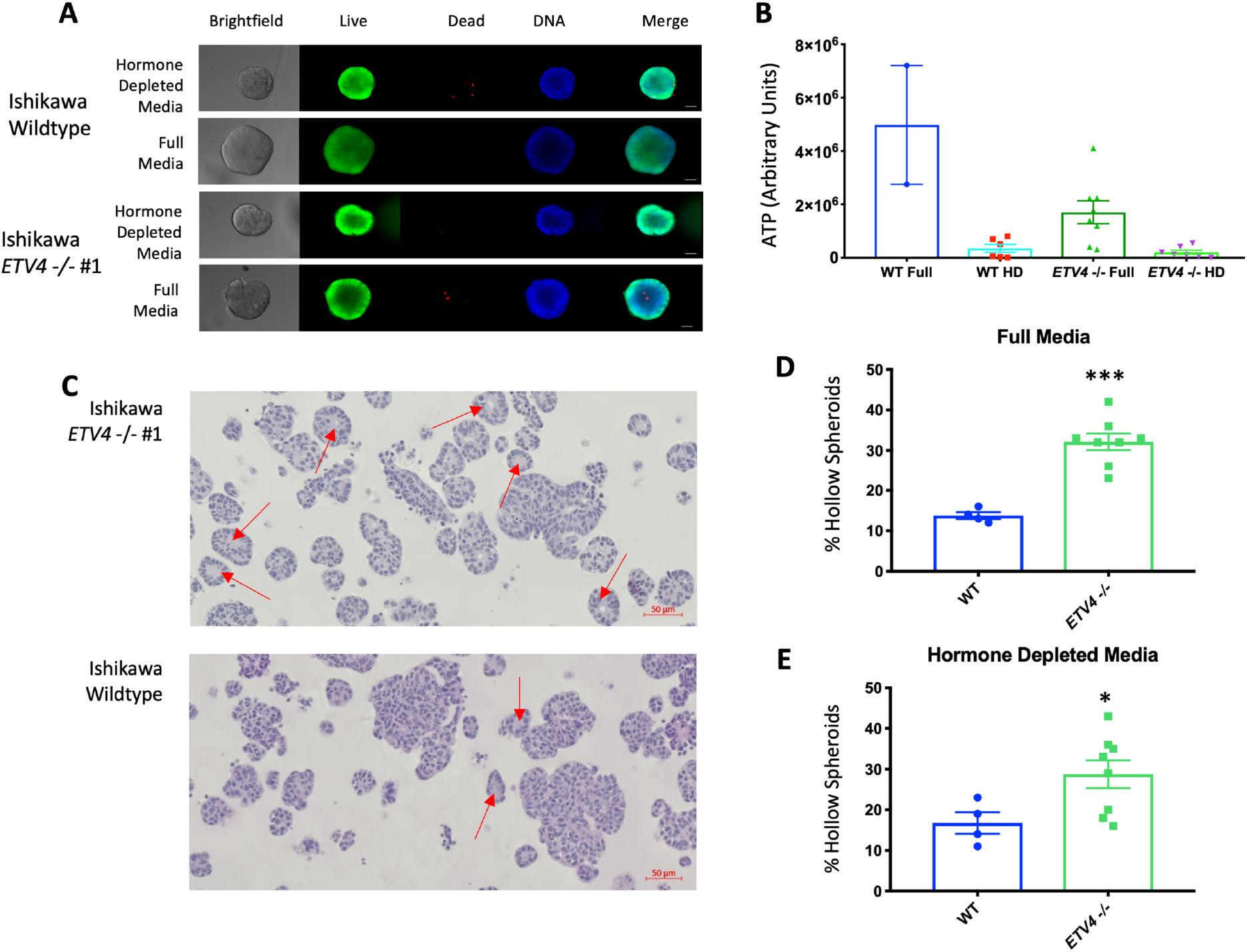
Loss of ETV4 leads to growth phenotypes. A) Representative images and staining of wildtype and *ETV4* KO organoids (Live cells - calcein AM; Dead cells - Cytox orange; DNA – Hoescht, scale bar=25μm). B) Loss of *ETV4* reduces metabolic activity in full serum media, but not in hormone-depleted (HD) media (WT Full (n=2), WT HD (n=6), *ETV4* -/- Full (n=8), *ETV4* -/- HD (n=7)). C) H&E staining depicts hollow organoid structures (red arrows) grown in full media, enriched in the knockout cells. Quantification of hollow organoids shows an increase in the knockout clones in both full serum media (p-value= 0.0001, unpaired t-test) (D) and hormone-depleted media (p-value=0.0474, unpaired t-test) (E) (WT Full (n=4), *ETV4* -/- Full (n=8), WT HD (n=4), *ETV4* -/- HD (n=8)). Error bars represent s.e.m.

### *In vivo* growth is reduced with *ETV4* loss

Based on the reduced growth of cells without *ETV4* in culture, we tested the ability of wildtype and *ETV4* knockout cells to grow *in vivo*. Cells were implanted into immunocompromised mice that had been ovariectomized and supplemented with or without an E2 pellet. We observed a 1.6-fold reduction in tumor growth (p-value = 0.0003, ANOVA) due to ETV4 loss in the no supplementation setting (Fig. 7A). In the presence of an E2 pellet, wildtype Ishikawa cells grew approximately 10-fold faster, indicating a strong stimulation of growth by E2. With E2 supplementation the growth difference between wildtype and *ETV4* knockout cells was more substantial (3.2-fold difference, p-value < 0.0001, ANOVA) (Fig. 7B). These results indicate that ETV4 is important for endometrial cancer cell growth *in vivo*, particularly in the context of estrogen stimulated growth.

**Figure 7.**
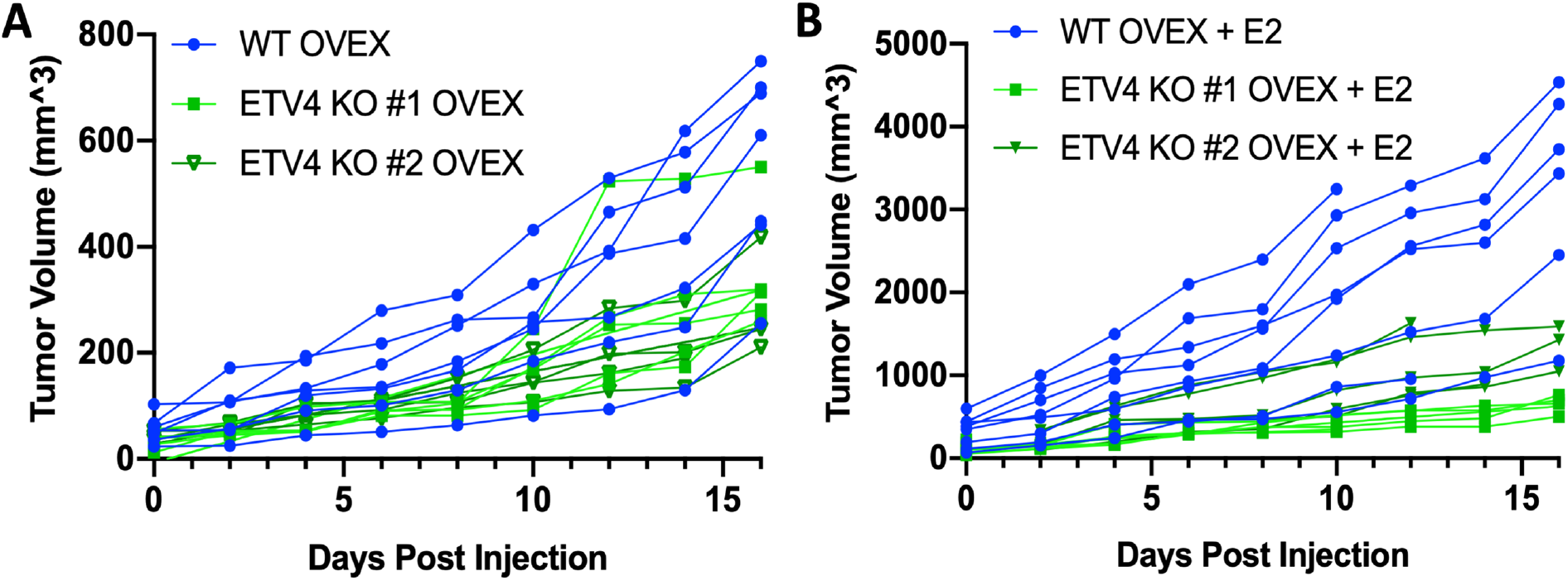
*ETV4* knockout cells exhibit reduced growth *in vivo*. Wildtype (blue, n = 16) and *ETV4* knockout cells (shades of green, n = 10 for each line) were implanted into ovariectomized immunocompromised mice. Half of each cohort was given E2 supplementation. Growth curves show tumor volume of wildtype and ETV4 knockout cells in mice without E2 supplementation (A) and with E2 supplementation (B). Day 0 is the first measurement day, which is 14 days after cell implantation.

## DISCUSSION

ER genomic binding is highly cell type-specific because it is dependent on cell type-specific transcription factors and chromatin accessibility. In breast cancer cells, FOXA1 and GATA3 play a critical role in determining and enabling ER genomic binding [19, 20]. However, in endometrial cancer cells, which exhibit different ER genomic binding patterns, the transcription factors that control ER binding site selection are unknown. In this study, we focused on ETV4, an ETS factor that exhibits overexpression specifically in endometrial cancer and its binding motif is enriched at endometrial cancer-specific ER binding sites.

To determine the importance of ETV4 in estrogen signaling, we used CRISPR/Cas9 to knockout *ETV4* in endometrial cancer cells. Genetic deletion of *ETV4* resulted in a large reduction of ER binding signal at the majority of bound loci across the genome. The level of reduction in ER binding is somewhat surprising because ETV4 overlaps with approximately half of ER binding sites, but *ETV4* knockout impacted approximately 70% of ER binding sites. Similar results were observed with *FOXA1* knockdown in breast cancer cells[19], suggesting that ER supporting transcription factors can impact ER binding at loci where their binding is not detected. The reduction that we observed was not due to changes in ER protein expression and must involve either ETV4 binding that is below ChIP-seq detection limits or secondary effects of ETV4 loss. Through the use of siRNA knockdown, we determined that the effects of ETV4 loss on ER genomic binding are immediate and do not require long-term gene expression changes. The large reduction in ER binding due to ETV4 loss resulted in an expected decrease in the transcriptional response to E2 treatment, indicating that loss of ER binding had transcriptional consequences.

Our original hypothesis for how ETV4 was impacting ER binding was that ETV4 is important in maintaining sites of open chromatin for ER. ATAC-seq analysis showed that some loci are impacted by ETV4 loss, but chromatin accessibility at the majority of ETV4 and ER bound loci are unaffected by *ETV4* knockout. The subtle changes in chromatin accessibility suggest that ETV4 is not acting as a pioneer factor for ER, unlike the pioneer factor role that FOXA1 plays for ER in breast cancer[19]. The lack of pioneer activity for ETV4 was supported by ectopic expression experiments in breast cancer cells. When we introduced ETV4 into T-47D cells, we found that ER binding site selection was unaffected, with possibly lower signal, and maintained a breast cancer binding profile. In addition, ETV4 binding was highly correlated with the chromatin accessibility pattern of T-47D cells. ETV4 does not appear to be playing a pioneering role in endometrial cancer cells and increased expression of ETV4 only aids in ER genomic binding in certain contexts.

ETV4 is impacting endometrial cancer ER binding to a similar level as FOXA1 in breast cancer, but the mechanism by which ETV4 is enabling ER binding is likely different than FOXA1’s pioneering role. Since we did not observe major changes in chromatin accessibility in absence of ETV4, we explored whether ETV4 might be affecting ER activation in another manner. We found that the ratio of nuclear to cytoplasmic ER was largely reduced by *ETV4* loss. This change in localization does not appear to be a general steroid hormone receptor defect in *ETV4* knockout cells, as no changes in localization were seen for GR, another steroid hormone receptor. How ETV4 is affecting ER’s localization remains unclear, though it may be doing this through multiple mechanisms. ER contains both an established nuclear localization signal and two regions considered non-canonical nuclear export signals[36]. Multiple post translational modifications have been shown to affect these sites, though it is not fully understood. The phosphorylation of Y537 has been indicated as necessary for ER interaction with exportin[37], and as ETV4 is present in the nucleus, it may play a role in preventing this phosphorylation event or other modifications, assisting ER in remaining in the nucleus. Another ETS factor, FLI1 has recently been shown to interact with GR, indicating a mechanism for ETS factors complexing with steroid hormone receptors[38]. While it remains unknown how ETV4 is affecting ER localization, this data suggests that ETV4 influences estrogen signaling at two levels: 1) ER activation and 2) chromatin accessibility maintenance.

The connection between ETS transcription factors and nuclear hormone receptors is intriguing as these gene families may have arisen together specifically in the metazoan lineage[39] and may have evolved to work together to control gene expression and downstream phenotypes. For example, ETV1 has been shown to reprogram androgen signaling, resulting in the formation of prostatic intraepithelial neoplasia[25]. ETS factors have been shown to be involved in regulating ER activity. The ETS factor ELF3 can bind to ER, blocking ER’s DNA binding domain and ability to regulate transcription[40]; however, this mechanism seems unlikely for ETV4, since ETV4 loss reduces ER activity. ETS factors can also promote estrogen signaling. For example, a breast cancer study found that ectopic expression of ETS1 led to an interaction with ER and promoted estrogen-dependent growth[41]. And a recent study found that ETV4 promotes estrogen induced proliferation and invasion in cholangiocarcinoma[42], although the mechanism is unknown. In agreement with these studies, we found that *ETV4* loss reduces growth of endometrial cancer cells. This effect was more pronounced in 3D culture and *ETV4* knockout spheroids formed hollow structures more readily than wildtype. While the implications of a hollow 3D growth pattern could be due to many factors, the resulting structures are similar to the glandular morphology observed in normal endometrium-derived organoids and are consistent with a less invasive and more differentiated cellular state [43]. However, we did not observe e-cadherin expression in the *ETV4* knockout organoids, indicating that full differentiation does not occur with ETV4 loss. We also observed a dramatic loss of ETV4 knockout cell growth *in vivo*. While some growth reduction was observed in the absence of estrogens, the growth differences between wildtype and ETV4 knockout cells was more pronounced in the context of estrogen supplementation. Overall, our results suggest that ETV4 is a critical factor in controlling and enabling estrogen signaling in endometrial cancer cells.

## Acknowledgements

This work was supported by NIH/NHGRI R00 HG006922 and NIH/NHGRI R01 HG008974 (to J.G.) and the Huntsman Cancer Institute. Research reported in this publication utilized the High-Throughput Genomics Shared Resource and the Preclinical Research Resource, which provided animal services, at the University of Utah and was supported by NIH/NCI award P30 CA042014. A.C.R. was supported by diversity supplement R00 HG006922S1. Imaging was performed at the Fluorescence Microscopy Core Facility, a part of the Health Sciences Cores at the University of Utah. Microscopy equipment was obtained using a NCRR Shared Equipment Grant # 1S10RR024761-01. We thank Shuba Narayanan and L. Charles Murtaugh for their experimental help and reagents. We thank K-T Varley, Simon Currie, Alana Welm and members of the Varley lab, Graves lab, Don Ayer’s lab, and Gertz lab for their research suggestions.

## Data availability

Raw and processed data is available at the Gene Expression Omnibus (GEO) under the following accessions: GSE129803 (ChIP-seq), GSE129804 (RNA-seq), and GSE129802 (ATAC-seq).

## MATERIALS AND METHODS

### Cell culture

Ishikawa cells (Sigma-Aldrich, 2014) and T-47D cells (ATCC) were grown in full media (RPMI-1640 (Gibco), 10% fetal bovine serum (Gibco), and 1% penicillin-streptomycin (Gibco)). Five days prior to induction, cells were moved to hormone-depleted media with phenol-red free RPMI-1640(Gibco), 10% charcoal-dextran treated fetal bovine serum (Sigma-Aldrich), and 1% penicillin-streptomycin (Gibco). Inductions involved treating cells with DMSO, as a vehicle control, or 10nM E2 (Sigma-Aldrich) for either 1 hour for ChIP-seq and ATAC-seq, or 8 hours for RNA-seq.

### Endometrial cancer patient-derived xenograft models

Animals were housed in the Comparative Medicine Center at the University of Utah under standard conditions. All animal procedures were conducted following approved Institutional Animal Care and Use Committee (IACUC) protocols. In brief, endometrial cancer patient specimens were collected in the operating room by the Biorepository and Molecular Pathology (BMP) Shared Resource and transported in ice-cold RPMI-1640 medium (ATCC) to the lab. Each tissue specimen was mechanically separated into multiple pieces using a sterile disposable scalpel (Andwin Scientific). A minimum of three 6-8 week-old, female nude mice (nu/nu) (Jackson Laboratories) were used for tumor implantation. Animals were anesthetized by intraperitoneal injection of Xylazine-Ketamine mixture at a dose of 0.1mg/kg and a subcutaneous analgesia, buprenophine (Reckitt Benckiser Pharmaceuticals), at a dose of 0.1mg/kg. A longitudinal 1 cm incision was made in the lower midline of the abdominal wall to penetrate the skin and peritoneum and then the uterus was exposed. A 2 mm incision was made on one horn of the mouse uterus. Tumor fragment (~4 mm^3^) was implanted into the mouse uterus under the microscope and the incision was closed using 6-0 prolene suture (Ethicon). The peritoneum and abdominal wall were sutured using 4-0 PERMA-HAND™ silk sutures and 4-0 polyglycolic acid sutures (Ethicon) respectively and then disinfected with alcohol/iodine. The nude mice were not given supplemental estrogen. E2 was monitored in the blood and found to be in the range of 25-50 pM; however, since the PDX were implanted in the orthotopic site, E2 levels may have been higher in the tumor environment. Tumor size was measured twice per week. Necropsy was performed once tumor size reached 2 cm in diameter. Clinical attributes can be found in Table S1. Models are available upon request.

### Generation of knockout lines

Ishikawa cells were grown in a 6-well plate in full media and co-transfected with the PEA3 (alias for ETV4) CRISPR/Cas9 KO plasmid (Santa Cruz, sc-422185) and the PEA3 HDR plasmid (Santa Cruz, sc-422185-HDR) using Lipofectamine 3000. Cells were treated with puromycin at 1μg/mL beginning 72 hours after transfection. RFP-positive single cells were sorted using fluorescence activated cell sorting into each well of a 96 well plate. Screening for knockout clones was performed by western blot and qPCR, as described below. Ishikawa wildtype and *ETV4* knockout cells were authenticated using STR marker analysis and found to be Mycoplasma negative (IDEXX, tested April 2019).

### Western blots

Ishikawa cell pellets were lysed via sonication at 40% amplitude for 10 seconds, 3 times with 10 second rests on an Active Motif EpiShear probe-in sonicator, in RIPA (1XPBS, 1% NP-40, 0.5% sodium deoxycholate, 0.1% SDS) supplemented with protease inhibitor (Pierce Protease and Phosphatase Inhibitor). Debris was removed by centrifugation at 14,000g for 5 minutes at 4°C and discarding the pellet. Protein quantification was performed using the DC Protein assay (Bio-Rad). 40μg of protein was run on NuPAGE 4-12% Bis-Tris protein gel (ThermoFisher Scientific) and transferred to PVDF membrane using the iBlot 2 system. Membranes were blocked in 5% nonfat milk solution for 1 hour at room temperature, and incubated overnight at 4°C in primary antibodies: ETV4 (Aviva ARP 32263_P050; 1:500 dilution), ERα (Santa Cruz HC-20; 1:1000 dilution), β-actin (Santa Cruz, sc-47778; 1:500 dilution), GR (Cell Signaling D6H2L; 1:1000 dilution), Lamin (Abcam 16048, 1:1500 dilution), or Tubulin (Molecular Probes, 236-10501; 1:20,000 dilution). Membranes were incubated for 1 hour at room temperature in appropriate secondary antibodies: Goat anti-mouse IgG (H+L) HRP (ThermoFisher Scientific) or Goat anti-rabbit IgG (H+L) HRP (ThermoFisher Scientific). Blots were developed using Super Signal West Femto Maximum Sensitivity Substrate (ThermoFisher Scientific).

### Immunohistochemistry

Tumors were sectioned by ARUP Research Histology services. Paraffin removal and rehydration of samples was performed with sequential steps in xylene (twice for 6 minutes), followed by 2 minutes in 100% ethanol (EtOH) (twice), 95% EtOH, 70% EtOH and 40% EtOH. Samples were then transferred to PBS. Peroxidase inactivation step was performed in 1% H_2_O_2_ for 20 minutes on shaker. Samples were washed in PBST 3 times for 3 minutes each. Antigen retrieval was performed using a 1:100 dilution of Vector Labs Antigen Unmasking solution in a PickCell 2100-Retriever; and samples were allowed to cool to room temperature. Samples were washed in PBST 3 times for 3 minutes each. Blocking was performed after marking label edge with PAP pen, using 400µl blocking solution (5% donkey serum in PBS + 3% Triton-X-100 + 0.2% sodium azide) in humid chamber for 1 hour at room temperature. Blocking solution was tapped off, replaced with 400ul of 1:100 dilution Millipore ETV4 (HPA005768) in blocking solution and incubated at 4°C overnight. Slides were washed 3 times for 3 minutes in PBST each and incubated in biotinylated rabbit antibody (Jackson Immunoresearch #711-065-152) 1:250 dilution in blocking solution for 1 hour at room temperature. Slides were washed 3 times for 3 minutes and incubated in Vector ABC Elite reagent (PK-6100) for 30 minutes at room temperature. Detection was performed using Vector DAB substrate (SK-4100). Counterstain was performed using hematoxylin. Slides were dehydrated and coverslips were added. Images were taken using an Olympus CX41 microscope and MicroSuite software.

### Generation of rescue lines

*ETV4* rescue experiments were performed by infecting cells with *ETV4* expressing retrovirus using a construct described previously[44]. Expression and infection of retrovirus was performed by standard protocol. In brief, 2mL of viral supernatant was added to a 50% confluent 100 mm dish of Ishikawa cells. 24 hours after infection, cells were selected for integration using 200μg/mL hygromycin, for a total of 14 days under selection. The rescue lines were validated by Western blot and qPCR looking at ETV4 expression.

### 3D culture

Cell lines were plated as 500 cells/well in 48-well plates (MatTek, 6mm, glass-bottomed), embedded in 9μL of phenol-red free matrigel matrix (Corning). Organoids were allowed to form for three weeks with media changes every 3-4 days. Live/dead imaging was performed by staining with Hoechst (5µg/mL), Calcein AM (1µM), and SYTOX Orange (0.5µM) for 1 hour prior to imaging with an Olympus IX81 inverted microscope. Relative ATP levels were measured using CellTiter Glo 3D (Promega). Fully-mature spheroids were formalin-fixed, paraffin embedded and sectioned. Sections were stained for hematoxylin and eosin (H&E) and imaged on a Zeiss Axio Scan.Z1. Quantification of hollow spheroids was done by a blind count of hollow spheroids by visual inspection from four equally sized regions of each H&E stained slide, analyzing only spheroids that were at least 50μm across.

### ChIP-seq

For Ishikawa cells, cells were grown as described above in hormone-depleted media in 15 cm dishes. After 1-hour treatment with E2 or DMSO, cells were crosslinked with 1% formaldehyde for 10 minutes at room temperature. Crosslinking was stopped with 125mM Glycine and cells were washed with cold PBS. For PDX tumors, 200mg of each sample was processed as previously described[45]. ChIP was done as described previously[46], with the following modifications. Sonication was performed at 40% amplitude, with 8 cycles of 30 seconds, with 30 seconds of rest after each cycle, on an Active Motif EpiShear probe-in sonicator. The antibodies used were: Santa Cruz ERα (HC-20) and Sigma FLAG M2 for ETV4; validation with Millipore ERα (06-935). Libraries were sequenced on an Illumina HiSeq 2500 as single-end 50 basepair reads. Reads were aligned by bowtie[47] to the hg19 build of the human genome using the parameters: -m 1 -t –best -q -S -l 32 -e 80 -n 2. Peak calling was performed using MACS2[48] with a p-value cutoff of 1e-10 and the mfold parameter constrained between 15 and 100. To determine read depth of peaks called, bedtools coverage[49] was used to calculate the counts between sets. For motif enrichment analysis, ER ChIP-seq sites were scanned for matches to ETV4 and FOXA1 motifs[14] using Patser[50]. To calculate fold enrichment, the number of significant motif matches was compared to the number of significant motif matches in randomly shuffled sequences that maintained nucleotide frequency.

### Quantitative PCR

Cells were harvested from 24-well plates in 350μL RLT Plus (Qiagen) supplemented with 1% beta-mercaptoethanol after an 8-hour induction with 10nM E2 or DMSO. Lysates were purified using the Quick-RNA Mini Prep kit (Zymo Research). Quantitative PCR was performed using the Power SYBR Green RNA-to-CT 1-Step Kit (ThermoFisher Scientific) using 50ng of total RNA for each sample with 1 minute of extension per cycle. Quantification was performed on a CFX Connect light cycler (Bio-Rad). Primers are listed in Table S2.

### RNA-seq

Cells were harvested from 10cm dishes in 600μL RLT Plus (Qiagen) supplemented with 1% beta-mercaptoethanol after an 8-hour induction with 10nM E2 or DMSO. RNA purification was performed using RNA Clean and Concentrator (Zymo Research) with the optional DNase I treatment. The KAPA Stranded mRNA-seq kit (Kapa Biosystems) was used to create Poly(A) selected libraries from 1μg total RNA. Reads were aligned to the human genome, hg19 build, using HISAT2[51]. Sam files were then converted to bam files and sorted via Samtools[52]. Read counts for each gene[53] were determined from alignments using featuresCounts from the SubRead package[54]. Normalization and differential expression analysis were performed using DeSeq2[55].

### ATAC-seq

In order to look at chromatin accessibility, 250,000 cells were collected after growth and a 1-hour E2 or DMSO induction in 10cm dishes. ATAC-seq was performed according to the protocol described in [32]. Libraries were sequenced on an Illumina HiSeq 2500 as single-end 50 basepair reads. Sequencing reads were aligned by bowtie[47] to the hg19 build of the human genome using the following parameters: -m 1 -t –best -q -S -l 32 -e 80 -n 2. Read counts were calculated using featureCounts from the SubRead package[54].

### Nuclear and cytoplasmic isolation

Cells were harvested from 15cm dishes after 1-hour inductions with 10nM E2, 100nM Dex, or DMSO. Harvest and isolation of nuclear and cytoplasmic fractions were performed using Nuclear Extract Kit (Active Motif). Samples were quantified and immunoblotted as described above. In order to compare nuclear amounts without oversaturation of signal, 4ug of nuclear fractions were run, alongside 30ug of cytoplasmic fractions.

### Knockdown of ETV4

Wildtype Ishikawa cells were grown in hormone-depleted media in 15cm plates and transfected with a non-targeting control siRNA or an siRNA that targets ETV4 (TriFECTa DsiRNA Kit hs.Ri.ETV4.13, Integrated DNA Technologies) at a final concentration of 10 nM using the Lipofectamine RNAiMAX transfection reagent (Thermo Fisher). Cells were harvested for protein, RNA and ChIP-seq at both 72 hour and 96-hour timepoints, followed by 1-hour inductions of 10nM E2 or DMSO. Knockdowns were validated by western blotting, quantitative PCR, and ChIP-seq as described above.

### In vivo studies

Eight to ten-week-old female NOD.Cg-*Prkdc^scid^ Il2rg^tm1Wjl^*/SzJ (NSG) mice (The Jackson Laboratory) were ovarectomized to prevent estrogen production, with half having an estrogen pellet inserted in the flank. Pellets are composed of beeswax and E2, with each 30mg pellet containing 1mg E2. Pellets were produced as described in DeRose et al. [56]. Mice were injected with 3×10^6^ cells (in 20ul matrigel pellets) subcutaneously into the right mammary fat pad. 16 mice were injected with wildtype Ishikawa cells, half with E2 pellets. 10 mice were injected with each *ETV4* -/- line, half with an E2 pellet. Caliper measurements were started 14 days after implantation and measurements were every two days, until mice were sacrificed due to tumor burden.

## Notes

https://www.ncbi.nlm.nih.gov/geo/query/acc.cgi?acc=GSE129805

