## Supplemental Figures and Tables for "ETV4 is necessary for estrogen signaling and growth in endometrial cancer cells"

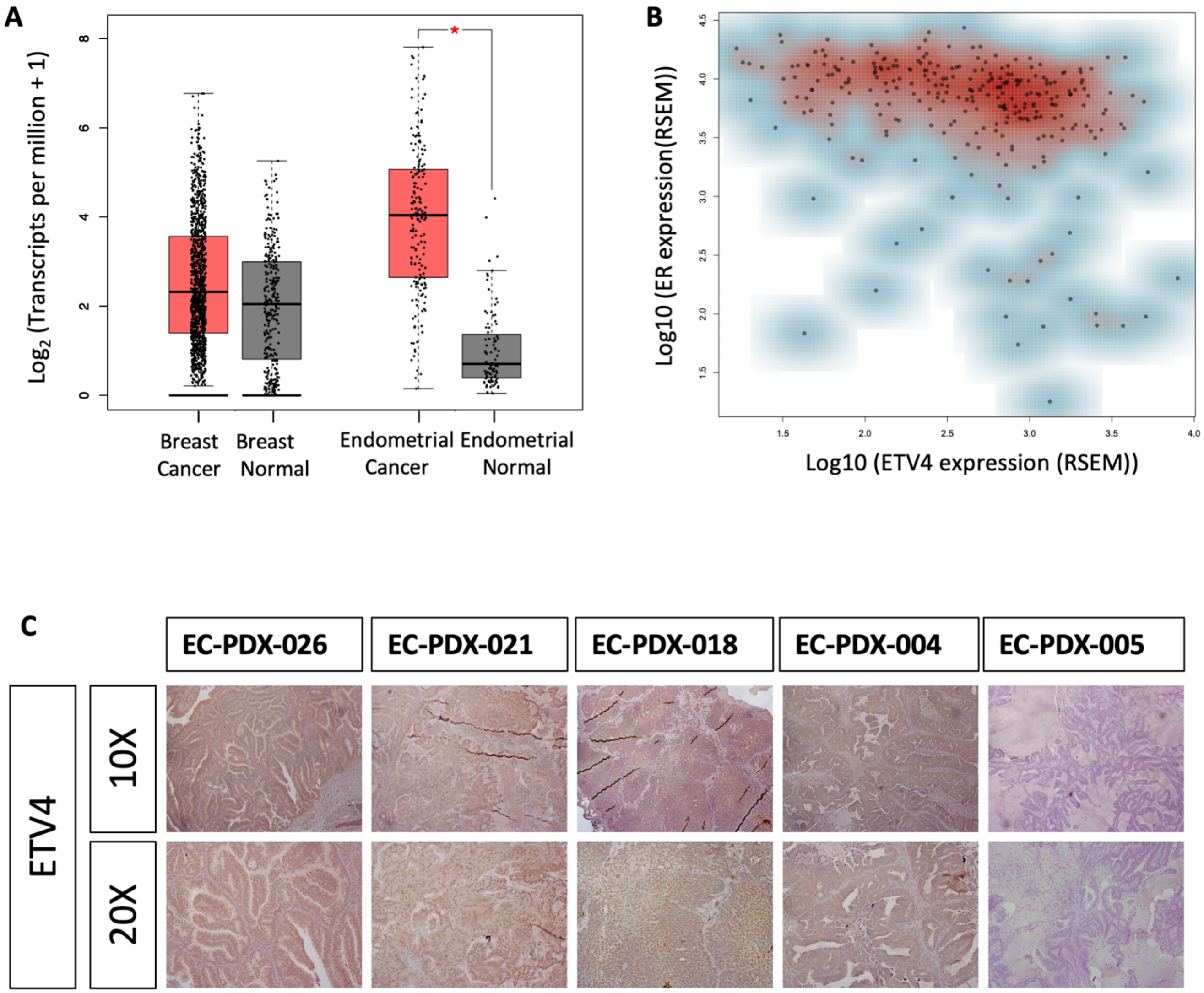


**Figure S1. ETV4 expression in endometrial cancer.** A) The expression of *ETV4* is shown in breast tumor and normal samples as well as endometrial tumor and normal samples from TCGA. There is no significant difference in *ETV4* expression between normal breast tissue and breast cancer, but a significant increase of expression in endometrial cancer compared to normal endometrium (p-value= 1x10-21). B) Plot shows the mRNA levels of *ETV4* and *ER* expression in endometrial cancer samples from TCGA. C) ETV4 expression in the nucleus is detected by immunohistochemistry (IHC) in three of five patient derived xenograft lines. Samples are ordered by ETV4 expression from left (highest) to right (lowest).

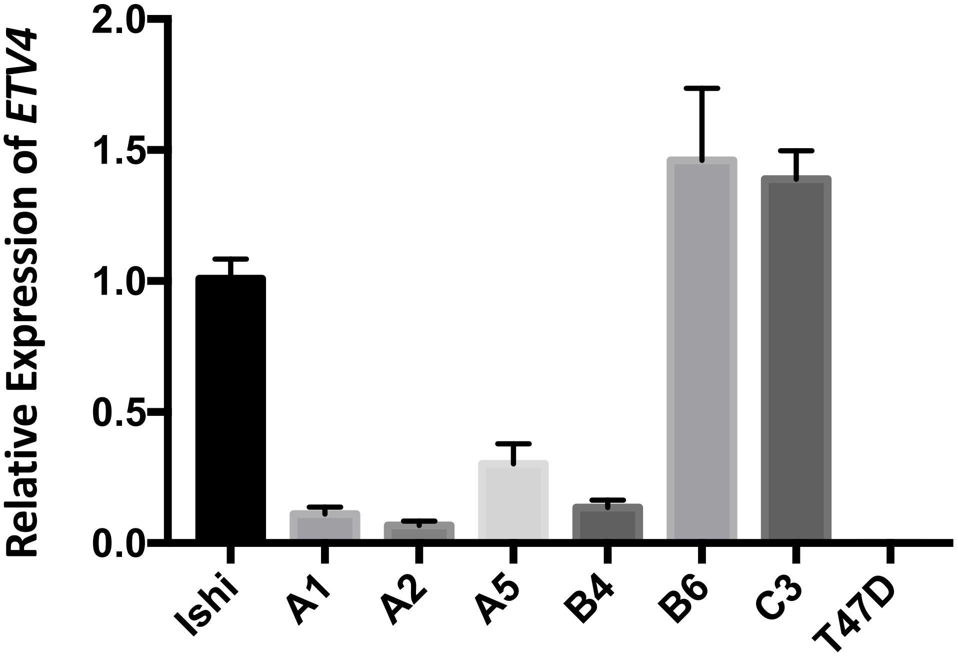


**Figure S2. Expression levels of *ETV4* in candidate knockout clones.** Quantitative PCR was used to measure mRNA levels, which were first normalized to CTCF and then normalized to wildtype expression levels. Clones A1 and A2 were selected as successful *ETV4* knockout lines, and used for all experiments. Error bars represent s.e.m.

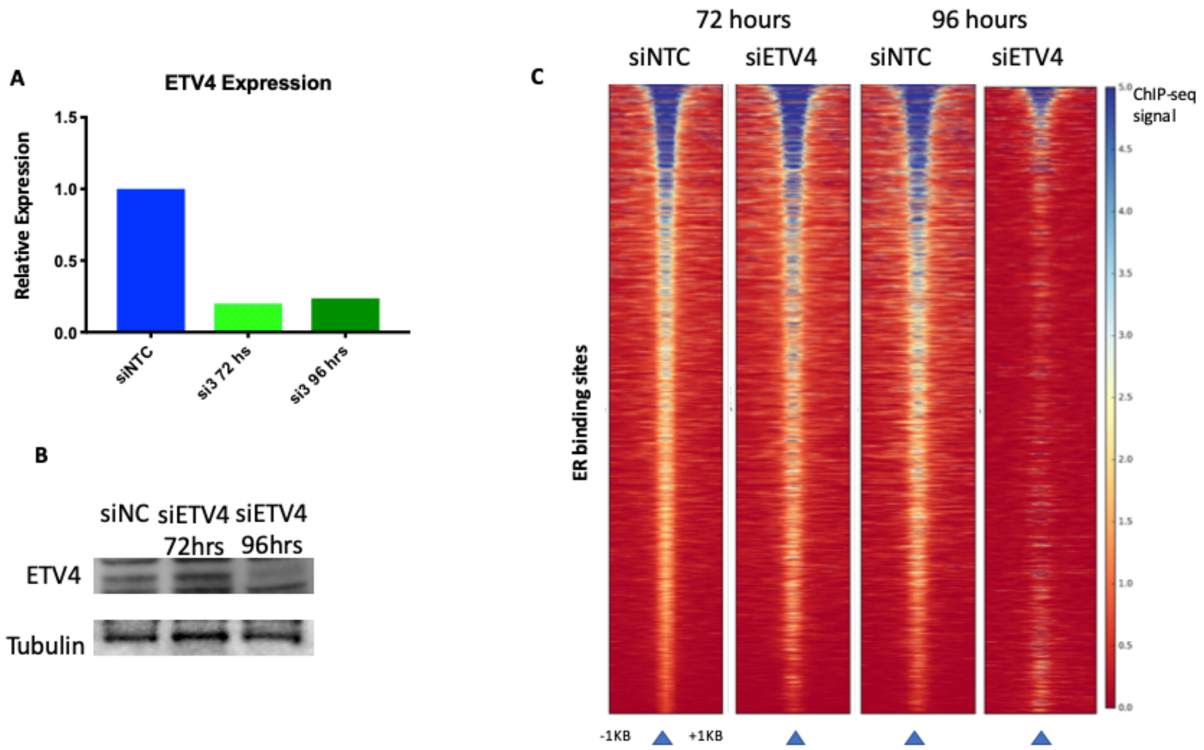


**Figure S3. Short-term knockdown of ETV4 reduces ER genomic binding.** A) Quantitative PCR and B) western blotting indicate successful knockdown of *ETV4* 96 hours after siRNA transfection. Tubulin is used as a loading control. C) Heatmaps of ER ChIP-seq experiments indicate successful reduction of ER binding with the transient loss of ETV4 at 96 hours. siNTC indicates non-targeting controls.

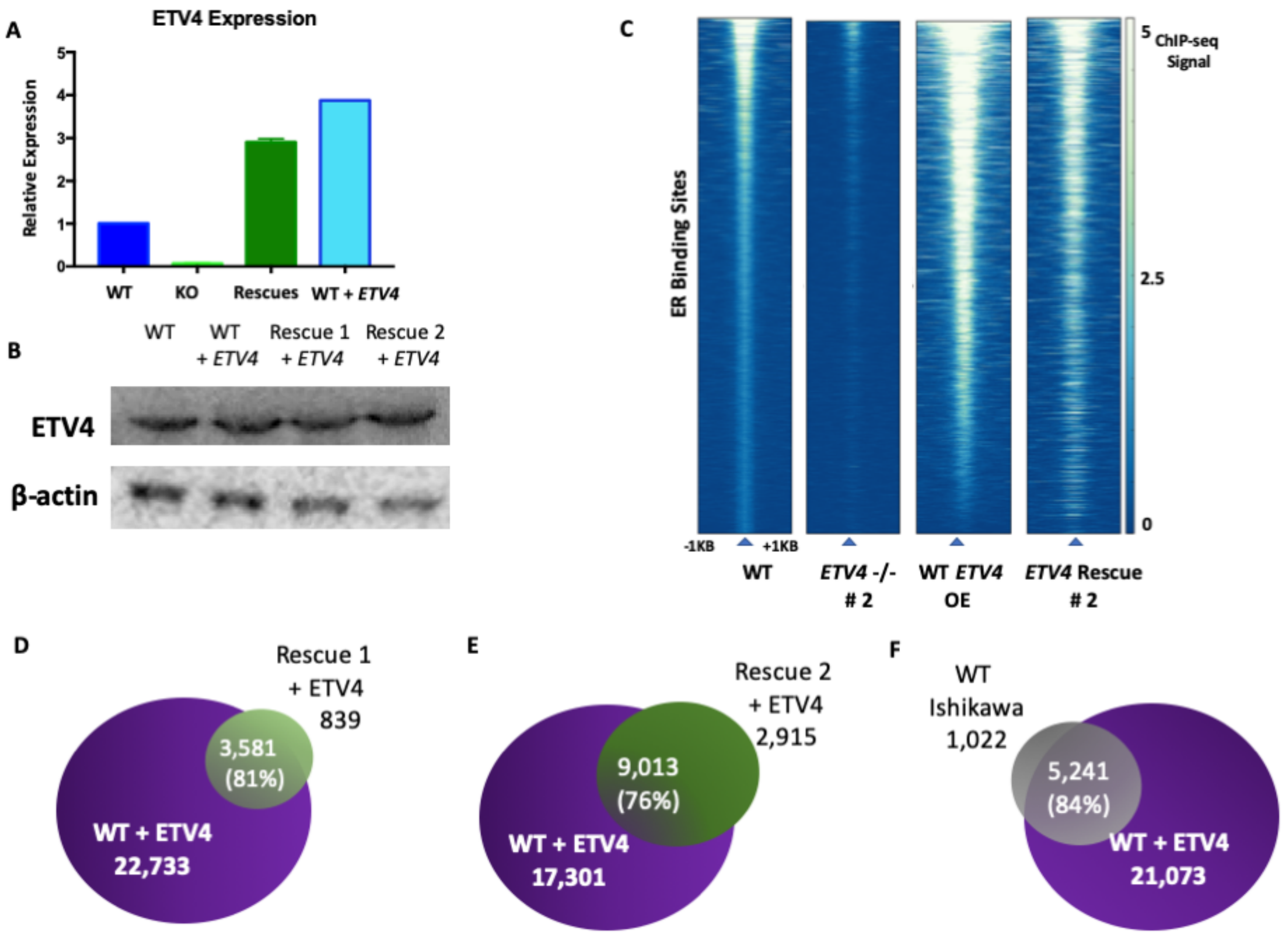


**Figure S4. Validation of rescue and overexpression of *ETV4*.** A) *ETV4* expression levels, measured by quantitative PCR and normalized to wildtype levels, indicate successful reintroduction of *ETV4* in knockout lines and overexpression in wildtype lines to levels above endogenous Ishikawa *ETV4* expression. B) Western blot indicates presence of ETV4 protein in rescue and overexpression lines. C) ER ChIP-seq signal after rescue of *ETV4* in knockout lines, alongside overexpression (OE) of *ETV4* in wildtype cells, compared to the original ER ChIP-seq shows a significant rescue of ER binding. ER binding sites in the rescue lines overlap by 81% (D) and 76% (E). 84% of ER binding sites in wildtype Ishikawa cells are found in Ishikawa overexpressing *ETV4* (F).

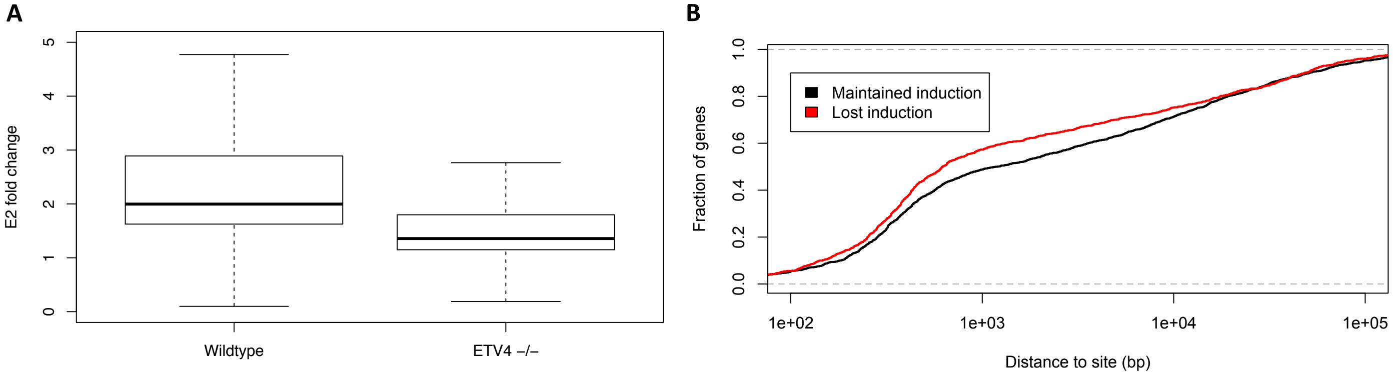


**Fig S5. Estrogen induced fold changes are higher in wildtype cells than *ETV4* knockout cells and genes that lose estrogen induction are closer to ETV4 bound sites.** A) Boxplot shows significant differences (p-value < $2.2x10^{-16}$, Wilcoxon test) in estrogen induced fold changes between wildtype cells and *ETV4* -/- as measured by RNA-seq. B) The cumulative distribution of distances between ETV4 bound sites and the transcription start sites of genes that loss (red) or maintain (black) an estrogen induction in ETV4 knockout cells.

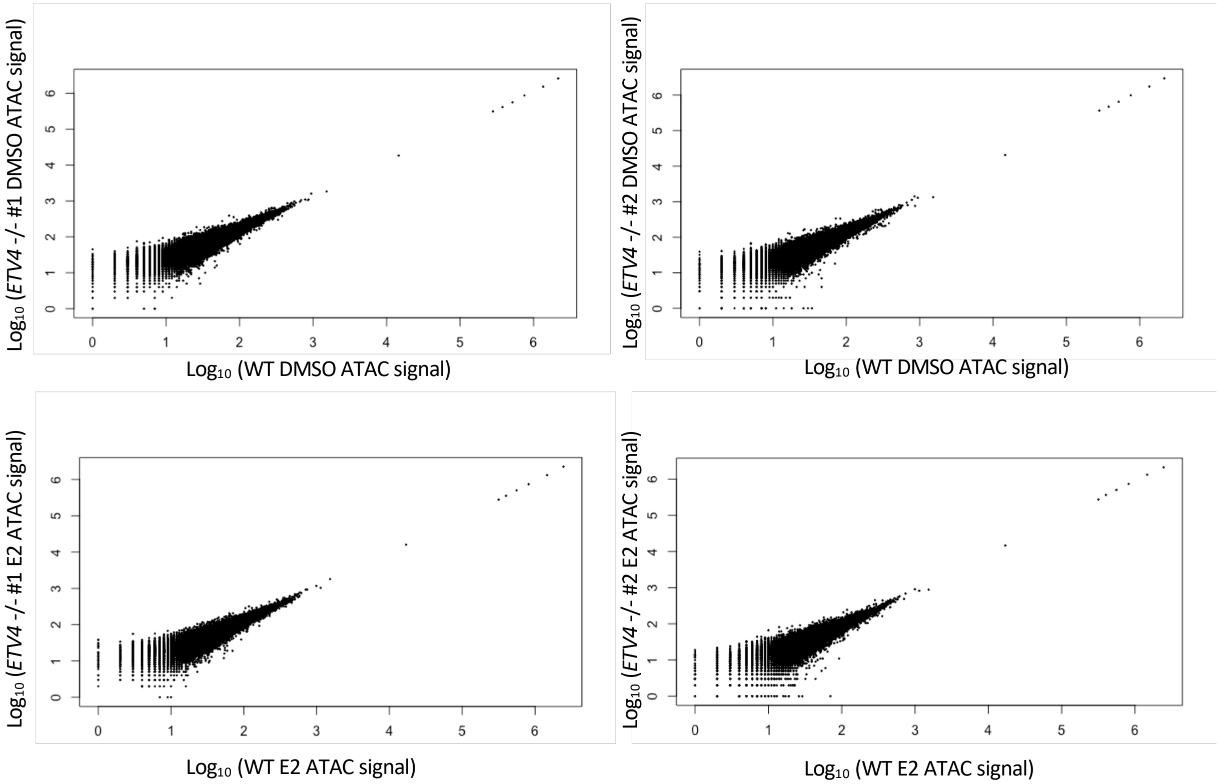


**Figure S6. Global chromatin accessibility is similar between wildtype and *ETV4* knockout lines.** Scatterplots compare the ATAC-seq signal at all identified accessible loci as reads per million in wildtype Ishikawa cells to *ETV4* knockout clones, after 1-hour DMSO (top panels) or E2 inductions (bottom panels).

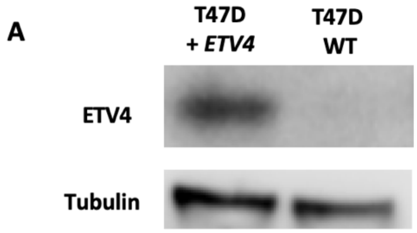


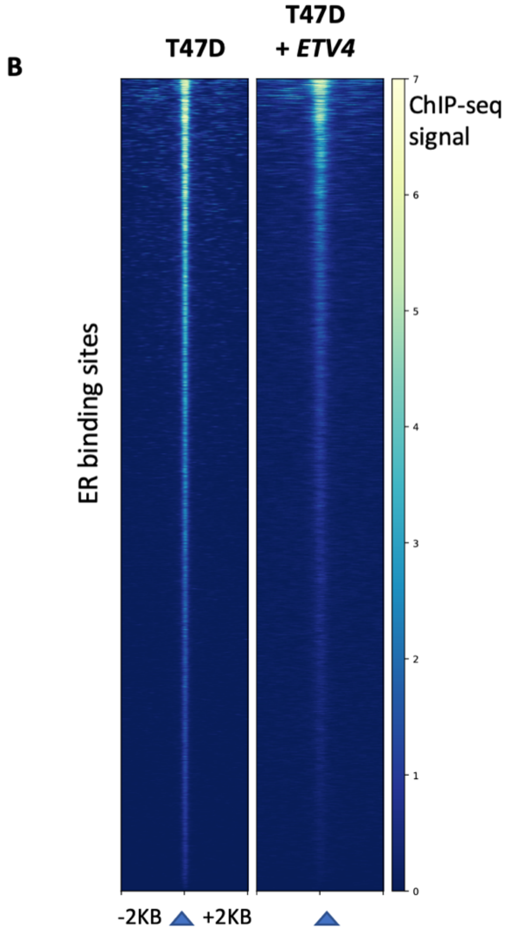


**Figure S7. Introduction of ETV4 in T-47D breast cancer cell line.** A) Western blot shows ectopic *ETV4* expression in T-47D cells. B) Heatmap shows similar ER ChIP-seq signal between T-47D cells and *ETV4* expressing T-47D cells.

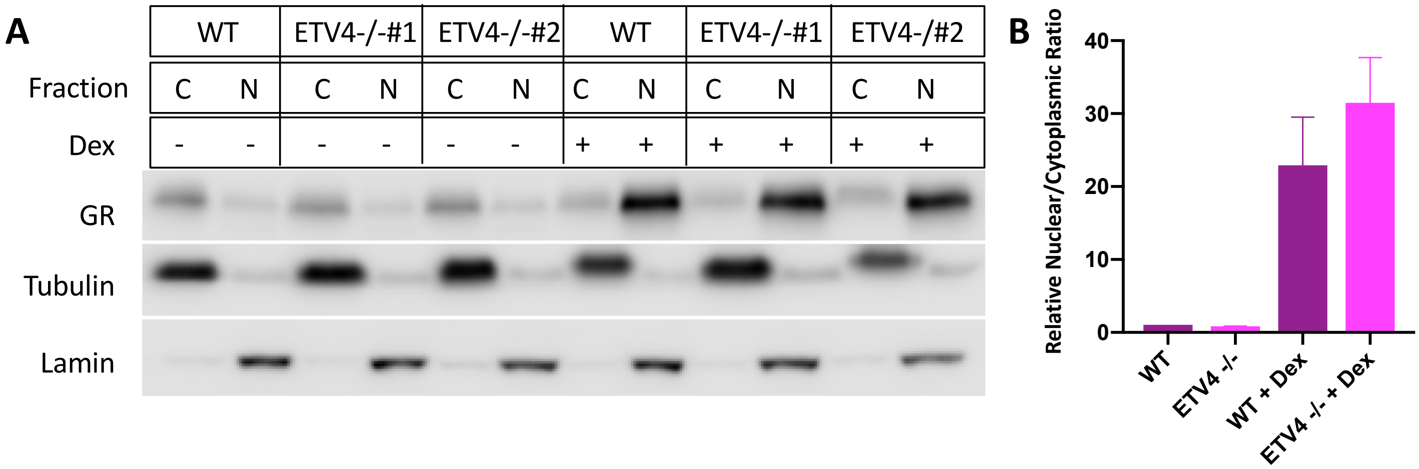


**Figure S8. GR localization does not change due to *ETV4* loss.** A) Western blot for GR in cytoplasmic and nuclear fractions of wildtype Ishikawa cells and both *ETV4* knockout clones. Tubulin was used as a cytoplasmic loading control and lamin as a nuclear loading control. B) Quantification confirms there is no change in the ratio of nuclear to cytoplasmic GR in ETV4 KO cells, with or without a 1-hour Dex induction.

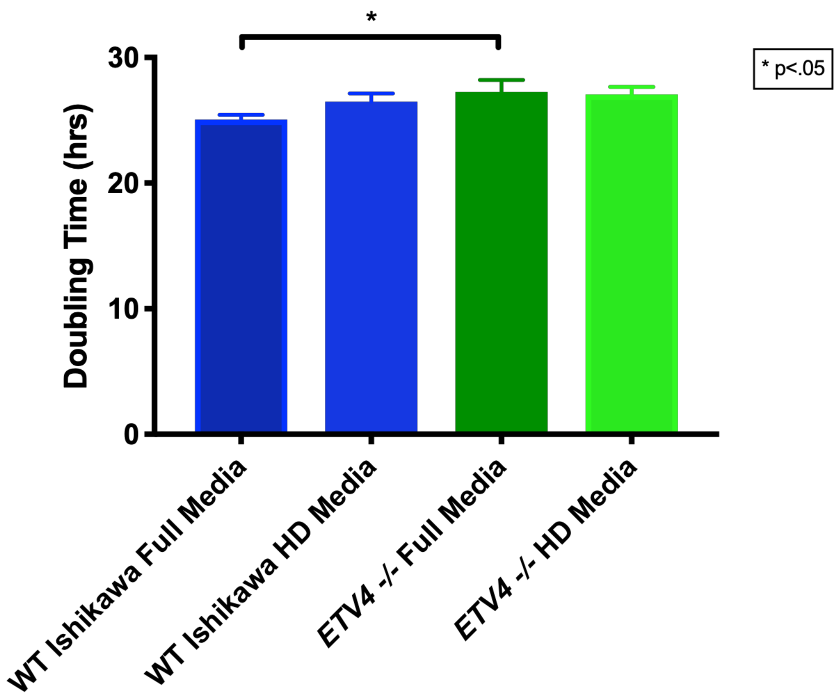


**Figure S9. Growth phenotype observed in 2D growth of *ETV4* knockout lines*.*** Confluency of cells grown in 2D culture was measured using live imaging on the IncuCyte Zoom, and doubling rates were calculated. The doubling rate of wildtype Ishikawa cells in full media (25.1 hours) is significantly faster than wildtype Ishikawa cells in hormone depleted (HD) media, (26.5 hours, p=.02, unpaired t-test). Loss of *ETV4* leads to an increase in doubling time in full media (27.3 hours, p=0.0012, unpaired t-test), but not in HD media (27.1 hours).

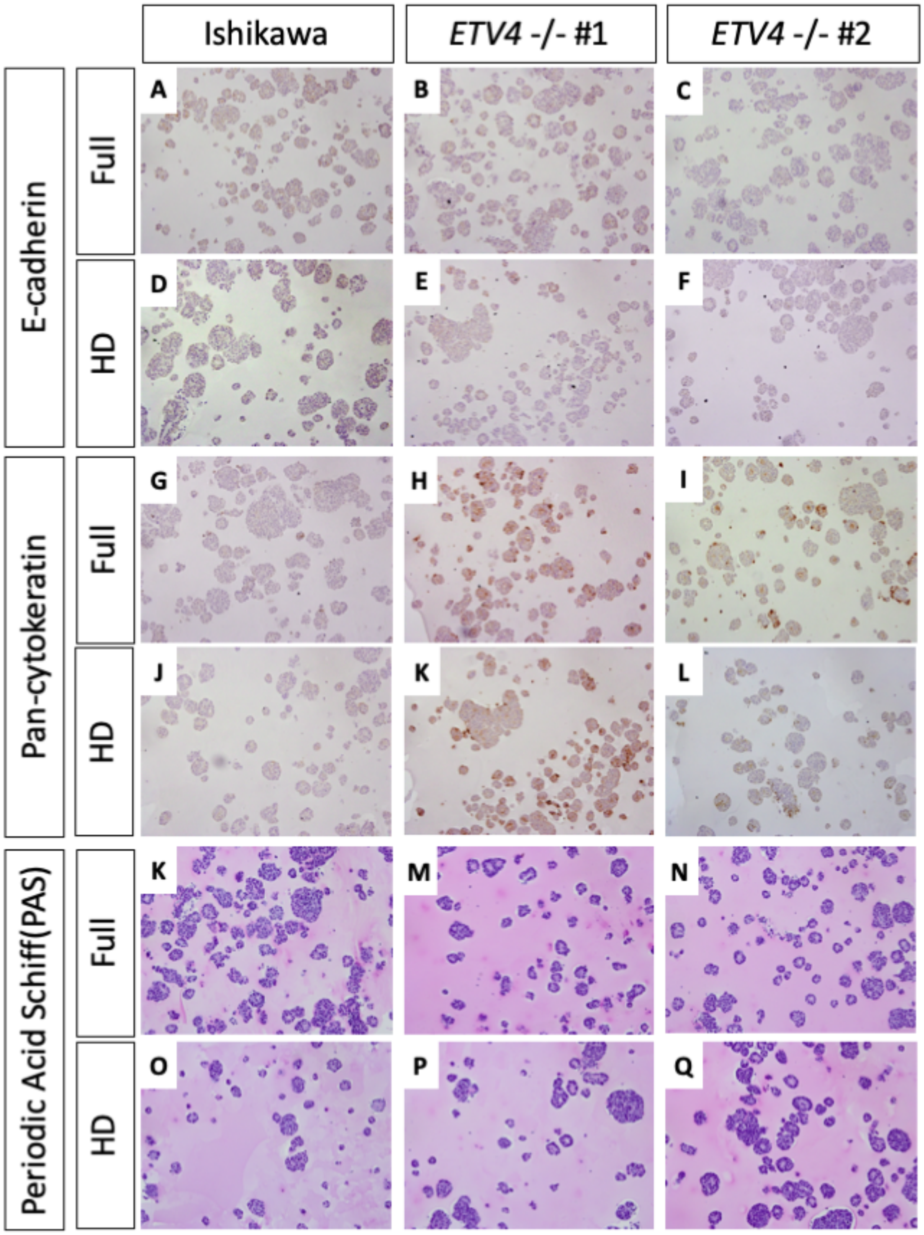


**Figure S10. *ETV4* knockouts increase expression of cytokeratins, a marker of normal endometrium.** E-cadherin IHC of wildtype Ishikawa and *ETV4* -/- organoids in full media (A-C) and hormone depleted (HD) media (D-F). Pan-cytokeratin IHC in full media (G-I) and HD media (J-L) indicating an upregulation in *ETV4* -/- lines. Periodic Acid Schiff mucin staining indicates an absence of mucins of all lines both in full media (K-N) and HD media (O-Q).

### Table S1. Endometrial cancer PDX clinical information

| Patient Information | | | | | | Xenograft Information | | | | |
|---|---|---|---|---|---|---|---|---|---|---|
| ID | Stage | Histology | FIGO grade | LN status | Metastasis | Grade | Metastatic site | Metastatic time | Median survival (days) | ETV4 staining |
| EC-PDX-004 | pT1a | EEA | Grade 1 | pN0 | None | Grade 1 | None | None | 187 | - |
| EC-PDX-005 | Relapse | EEA | Grade 2 | - | Peritoneum | Grade 2 | Peritoneum | 60 | 71 | - |
| EC-PDX-018 | pT1a | EEA | Grade 2 | pN0 | None | Grade 2 | None | None | 168 | ++ |
| EC-PDX-021 | pT1b | EEA | Grade 2 | pN0 | None | Grade 2 | None | None | 152 | ++ |
| EC-PDX-026 | pT1a | EEA | Grade 1 | pN0 | None | Grade 1 | None | None | 168 | +++ |

EEA = Endometrioid endometrial andenocarcinoma

LN = lymph node

### Table S2. *ETV4* knockout PCR validation primers

| Primer name | Sequence |
|---|---|
| *ETV4_qPCR_F* | AGGCGGAAACCTCGCCATCG |
| *ETV4_qPCR R* | CCAGAACCTCTAAGGTTTGC |
| *CTCF_qPCR_F* | ACCTGTTCCTGTGACTGTACC |
| *CTCF_qPCR_R* | ATGGGTTCACTTTCCGCAAGG |
